# Different dimensions of robustness - noise, topology and rates - are nearly independent in chemical switches

**DOI:** 10.1101/2020.08.01.232231

**Authors:** Sahil Moza, Upinder S. Bhalla

**Affiliations:** National Centre for Biological Sciences, Bengaluru

**Keywords:** bistable, chemical reaction network, robustness, motifs, evoltion

## Abstract

Life prospers despite adverse conditions in many unpredictable dimensions. This requires that cellular processes work reliably, that is they are robust against many kinds of perturbations. For example, a cellular decision to differentiate should be stable despite changes in metabolic conditions and stochasticity due to thermal noise. For evolutionary stability, the same differentiation switch should function despite mutations or the evolution of further regulatory inputs. We asked how cellular decision making responds to these three forms of perturbation, expressed in chemical terms as rate parameters, stochasticity, and reaction topology. Remarkably, we found that there was no correlation between noise robustness and either of the others and only a weak correlation between robustness to parameters and topology. Thus, a given chemical switch could be robust to noise yet sensitive to parametric or topological changes. However, we found families of reaction topologies derived from a common core bistable with symmetric feedback loops, which retained bistability despite the removal of reactions or substantially changing parameters. We propose that evolution involving chemical switches must navigate a complex landscape involving multiple forms of robustness, and the only way for a given switch to have a systematic advantage in robustness is to come from a ‘good family’ of mirrorsymmetric topologies.

**Significance Statement:** Life endures despite metabolic fluctuations and environmental assaults. For the thousands of cellular decisions to continue to work, they must be robust to these perturbations. Many cellular decisions are made and stored by chemical switches, which like light switches retain their state – on or off – even after the trigger is gone. We computationally explored what makes chemical switches robust. It turns out that some are robust to thermal noise, others to mutations that disable part of the switch, or to changes in chemical conditions. Surprisingly, these different forms of robustness are mostly independent. However, chemical switches come in families built around core reactions, and these families tend to score high or low on several measures of robustness.

## Introduction

Biological systems are adept at making decisions and remembering choices in the face of cellular and environmental change. Both decision making and memory formation are typically implemented in the cell using biochemical signaling systems that form bistable (YES/NO) switches, which encode each state as the local concentration of reactants. These decisions must be robust as they may determine the developmental fate of a cell (1, 2), or the state of synapses that encode a memory (3–5). How do such switches retain their state information reliably, despite the ongoing barrage of environmental, metabolic, and stochastic fluctuations?

The scale-free property of some biological networks, such as the postsynaptic proteome and metabolic networks grants them global robustness from random removal of nodes (6). However, if complex network are to be understood as intertwined networks of modular computations performed by motifs, local computations must be protected by locally robust networks. Indeed, there is evidence that local structural robustness is a good predictor of motif abundance (7), and the connections between motifs promote their stability (8).

There are two contradictory intuitions relating to survival and robustness. The first is that there is a tradeoff between different forms of robustness. This implies that you can select for, say, environmental resilience, but at the expense of evolvability. The other intuition is that if you have a system that works well over many environmental conditions, it will also work well if subjected to other perturbations like chemical noise. In either case, if one quantifies robustness along these different dimensions, one should see correlations. To look for these, one needs a large dataset and a crisp, quantifiable definition of robustness along each dimension. Bistable chemical switches meet these criteria (9, 10). While it is not easy to define all possible forms of survival challenge in chemical terms, there are at least three major dimensions where one can do so: mutations (signaling topology); thermal and metabolic environment (chemical rates); and noise (chemical stochasticity). We ask: 1, are bistable systems that are robust to one kind of perturbation also robust to other kinds of perturbations? 2, are certain chemical pathway structures more able to retain not just bistability, but robustness, as their topology evolves?

From an evolutionary perspective, molecular signaling networks are subject to continual change through mutations and gene rearrangements that layer additional regulatory mechanisms on top of existing pathways (11). A previous exhaustive examination of bistable switches has shown that they occur in families, with a simple root bistable topology that remains at the core of increasingly complex pathways (9). At an even more fundamental evolutionary level, bistable systems have also been proposed as chemical mechanisms for the homochirality observed in biological systems (12). Chirality is the property of molecules to be non-superimposable on their mirror reflections, called L and D forms. Most non-biological reactions produce molecules that are either achiral (superimposable) or heterochiral with an equal fraction of L and D enantiomer concentrations. However, biomolecules such as nucleic and amino acids are almost exclusively homochiral. Small bistable CRNs have been proposed as a mechanism to create homochiral products from achiral substrates (13, 14). Thus, a systematic comparison of several small bistable CRNs in the context of their robustness against various environmental perturbations may reveal their potential evolutionary success.

Bistability requires the presence of a core positive feedback loop (15), and other connections between reactants can alter the stability and robustness of these systems (16). Complex biochemical reaction networks such as signaling pathways, are exposed to different kinds of perturbations and have evolved to sustain their dynamical behavior (17). Such chemical systems are called *robust*, defined as the ability of a system to maintain its functions despite external and internal perturbations (11). Thus, the diversity of architectures across different biological systems may reflect the *robustness* of the underlying chemical network to the unique set of perturbations that the system may have encountered in its evolutionary past. For example, dendritic spines are very small, so their signal flow must work despite high levels of chemical noise (18–20). Similarly, the genome of a single-stranded RNA virus mutates rapidly, so its replication decision points must withstand possible topological change.

Here, we exhaustively analyzed all bistable reaction systems from a biologically-biased survey of chemistries possible with up to 4 reacting entities and 6 reactions (9). Then, we explored the robustness of these small bistable CRNs to changes in the reaction network structure (topology), parameters, and intrinsic noise to probe for relationships between different forms of robustness (21, 22). We found that there were some correlations between different forms of robustness over the entire dataset. Additionally, there were ‘family trees’ of topologies that contained a mirror-symmetric reaction topology involving autocatalytic feedback, which had higher parametric robustness. We propose that the dispersed landscape for multiple forms of robustness, combined with the family benefits of certain base chemical topologies, may be relevant to evolutionary trajectories of bistable dynamics.

## Results

We extracted 3561 bistable CRNs from the SWITCHES database and analyzed their chemical topology. All CRNs were constructed using reactions shown in Fig 1**A-C**. The smallest bistable CRN in the SWITCHES database is shown in Fig 1**D**. In Fig 1**E**, we demonstrate bistability using a 2D attractor plot of the chemical reaction system in Fig 1**D**, showing the 3 fixed points and the trajectories taken when started from different initial conditions.

**Fig 1.**
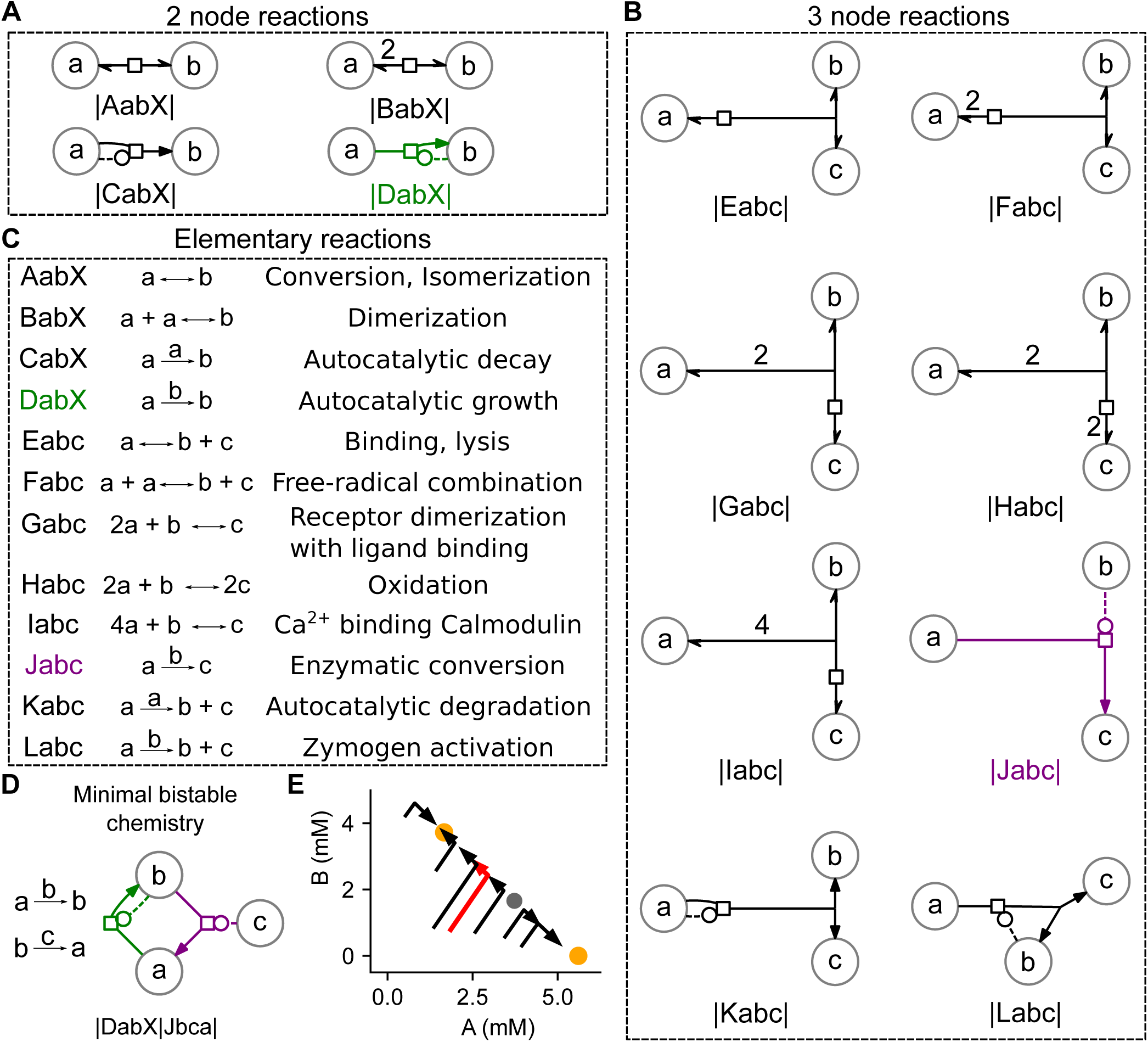
Constructing a bistable chemical reaction network from elementary reactions. **A, B)** Elementary chemical reactions in SBGN format with two and three participating reactants. Circles represent reacting entities and arrows represent reactions. Reaction signatures below in the CSPACE format. Each reaction is flanked by ‘|'. The capitalized first letter denotes 1/12 reaction types (A, B,… L). The next 2 or 3 letters are the reactant labels for reactions with 2 (**A**) and 3 molecules (**B**). **C)** Reactions graphically depicted in **A, B** represented in the canonical form with the reaction label (left). (Adapted from (9)) **D)** The smallest bistable CRN colored by the reaction types (**A**, **B**). **E)** Phase portrait of the bistable system in **C**, showing the two stable fixed points in orange and a saddle point in gray. Trajectories from several initial conditions are shown in black with one highlighted in red seen moving towards a fixed point and away from the saddle. For reaction DabX, K_m_ = 0.25 mM , k_cat_ = 6.97e-2 s^-1^ ; for reaction Jbca, K_m_ = 4.28e-06 mM, k_cat_ = 9.98e-03 s^-1^.

We then challenged all bistable CRNs with different forms of perturbations in order to calculate robustness (Fig 2**A**, 3**A**, 4**A**). First, we assigned randomized reaction rates to the CRNs within a physiological range and calculated the allowable variation in reaction rates while still retaining bistability (*parametric robustness*). Second, we calculated the amount of time a bistable CRN remains in one stable state before thermal fluctuations cause a state transition (*noise robustness*). Third, we systematically removed reactions from the CRNs to estimate the *structural robustness* of the CRN. Here we discovered CRNs as compositions of certain irreducible bistable CRNs called root CRNs. In the final section, we show the comparison of the different forms of robustness across all CRNs, and in the context of the composition of CRNs from different root CRNs.

**Fig 2.**
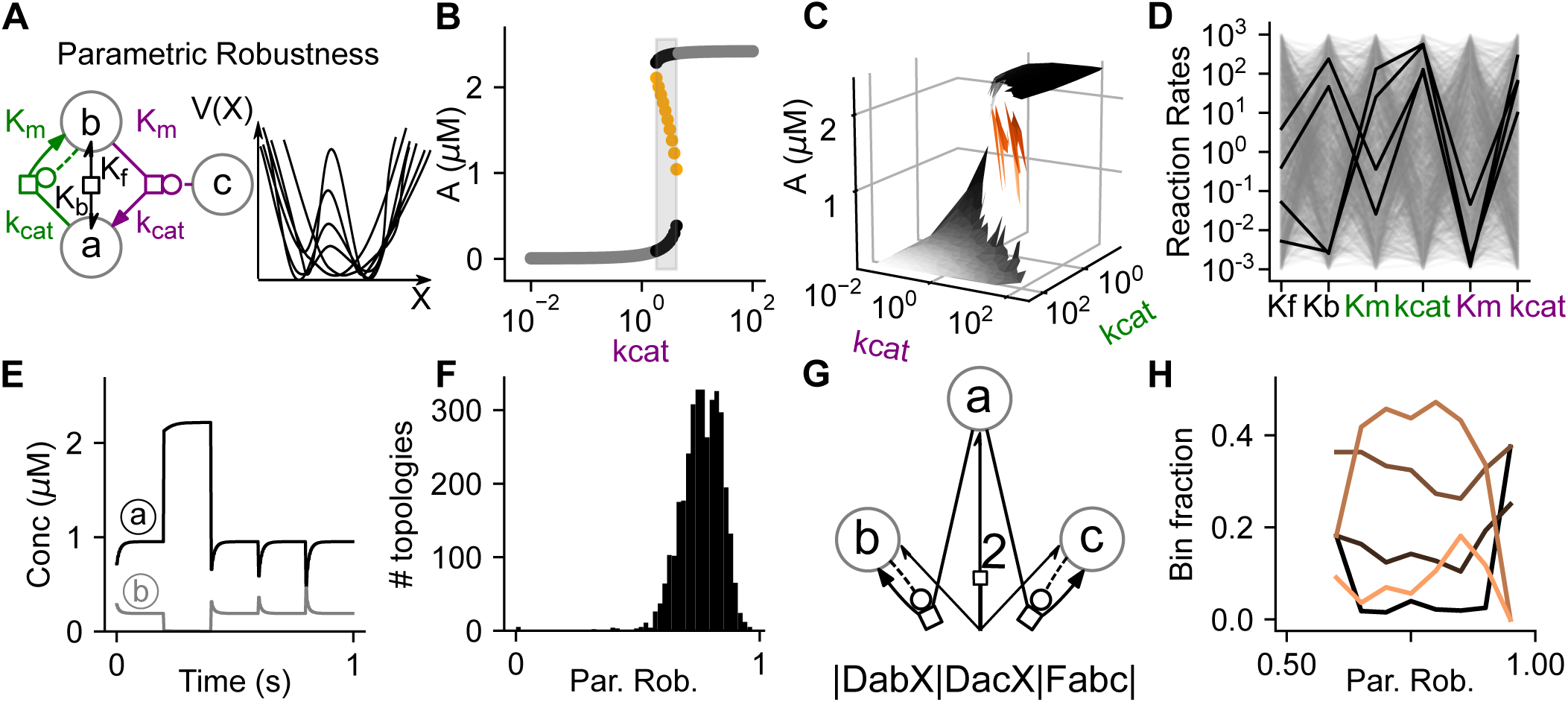
Parametric robustness in bistable chemical reaction networks. **A.** Schematic showing representative CRN and the changes in the basins of the fixed points due to change in reaction-rate parameters. **B.** Bifurcation diagram showing the region of bistability as a gray box, which appears in a narrow regime of parameter kc_at_ for the reaction marked in purple in Fig **2A. C.** Bifurcation diagram across two-different reaction-rates k_cat_ of green and purple reactions marked in **A**. **D**. All rate combinations sampled for the reaction topology marked in **A**. Dark lines indicate the combinations which were bistable. **E.** Deterministic simulation for one chosen bistable parameter set for the topology in **A**. Black and gray represent molecules a and b. The system was perturbed 4 times, and a state transition happened on each of the first two perturbations. **F.** The distribution of parametric robustness for all 3561 bistable topologies. **G.** The topology with the highest parametric robustness. **H.** Composition of topologies at different levels of parametric robustness. For different levels of parameter robustness (binned on x-axis), the proportions of topologies with different degrees of symmetry are shown in different colors. The orange lines denote the least symmetric CRNs and progressively darker lines denote increasing symmetry.

### Highly parametrically robust systems have a mirror-symmetric architecture

Parametric robustness for a bistable CRN is defined as the fraction of the sampled parameter space of chemical reaction rates that retains bistability (9). CRNs are employed as signaling cascades for transmitting information and performing various computations. In this sense, bistable systems carry binary codes within their steady-state concentrations. These steady-state concentrations change with reaction rates, and most CRNs are bistable only within a certain range of parameters.

We assigned 2000 randomly-sampled reaction rates within the biologically plausible range (Methods) to each of the 3561 CRNs, using the Latin-hypercube method for high-dimensional logarithmic sampling (23, 24). Each CRN instance with an associated set of reaction rates was called a *model*. We then tested these chemical kinetic models for bistability (Methods). We found that 531313 out of the total 7.122 million models (7.46%) were bistable, which is a rather large fraction considering the high dimensional (4 to 12 dimensions) parameter space from which we sampled. To obtain such a high fraction of bistable models, bistability would have to persist over roughly 4.5 log-10 units of the 6 log units we sampled for each parameter.

On further investigating the CRNs with high parametric robustness, we found that topologies that were highly robust to parameter variations also had symmetric chemical reaction diagrams (quantified below). The top parametrically robust topology is shown in Fig 2**G**, where the mirror-symmetric core of the network across the molecule *a* can be observed. This topology and many other parametrically robust CRNs had a “symmetric winner-takes-all” strategy. In Fig 2**G**, molecules *b* and *c* compete with each other by auto-catalyzing their own formation, with structurally identical interactions with *a*. If there is a slight excess of one over the other, the CRN is driven to a state with all molecules of one type and none of the other.

Next, we quantified the symmetry across all possible bistable CRNs. We calculated the size of the largest subnetwork of the CRN for which permutations of the node labels preserved the relationship between the nodes (Methods). We then calculated the *local symmetry* by quantifying the size of the asymmetric network for the CRNs where only a part of the network was symmetric. For this, we picked the globally symmetric topologies and for each, counted the number of extra reactions excluding the symmetric subnetworks (Methods). Using this approach, we grouped all topologies by the number of excess reactions, called the “order” of the topology. We found that the distributions of topologies that contained locally symmetric subnetworks were preferentially higher in parametric robustness than a random sample from the distribution (Fig 2**H**, 2-sample KS-test, p-value = 5e-07, 4e-10, 1e-50, 8e-56 for orders 1-4).

In summary, symmetric mutually-competitive autocatalytic reaction networks had the highest parametric robustness among all CRNs.

### Noise Robustness

Biochemical reaction networks are often compartmentalized in small volumes, such as dendritic spines, which undergo large signaling noise due to small copy numbers of reactants (25). For bistable systems, intrinsic (thermal) noise can lead to spontaneous state transitions between the two stable fixed points, which implies a loss of memory, imprecise decisions, and general loss-of-function (18, 25, 26).

Noise Robustness (*N*) at a given volume, was defined as the geometric mean of the average time spent in the two bistable steady-states before a spontaneous transition (*residence time*), averaged over all the different parameter combinations for that CRN. In Fig 3**B-D** we show the stochastic time-series evolution of molecule *a* from the model in Fig 3**A**, over 3 different volumes sampled logarithmically. In Fig 3**C**, for example, the concentration of molecule *a* stochastically transitions between the two different fixed points. We ran the stochastic simulation for each parameter set that was found to be bistable for each CRN for a period of 86400s (1 day) for 10 trials each.

**Fig 3.**
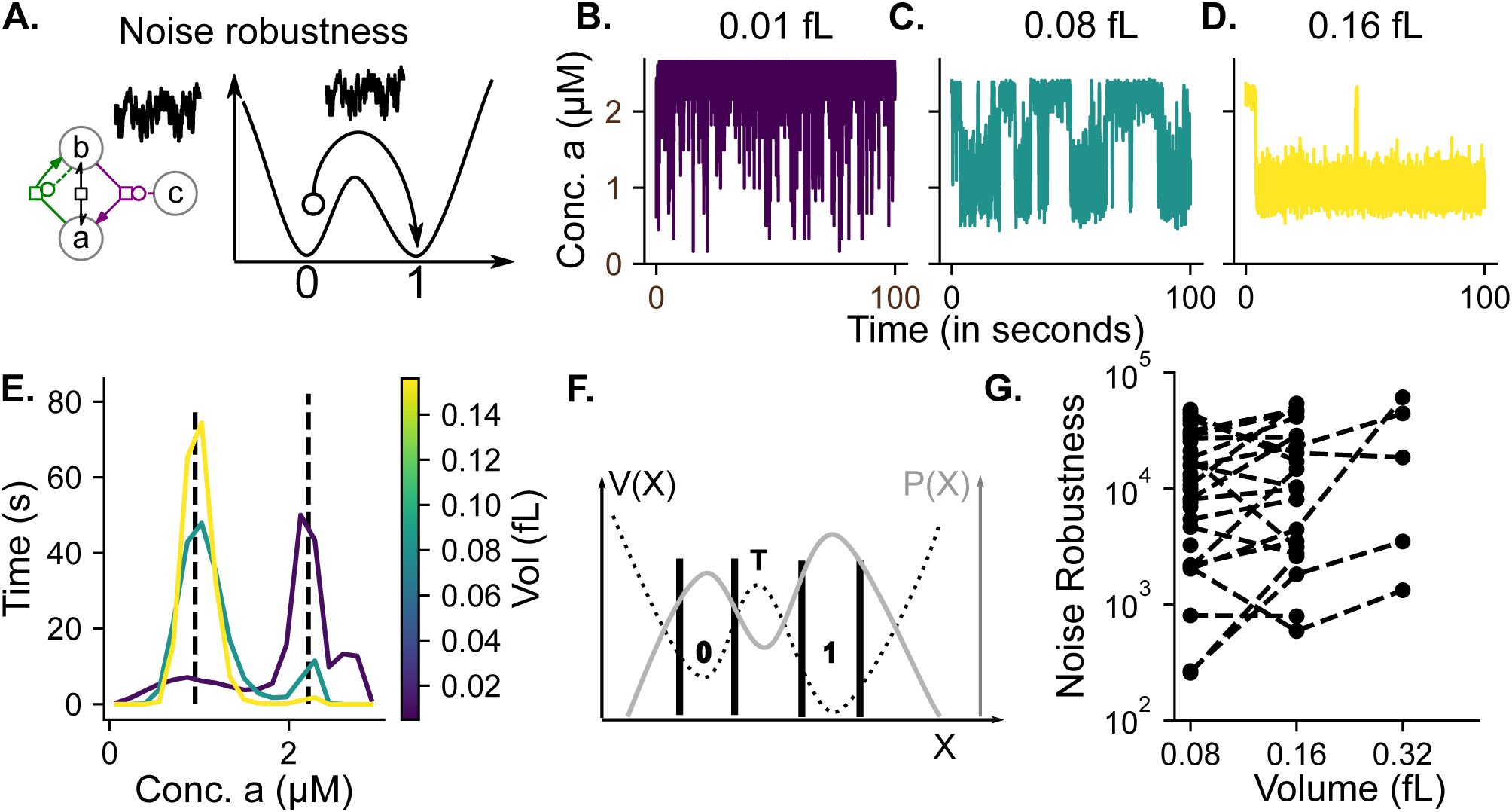
Noise robustness in bistable chemical reaction networks. **A.** Representative CRN and a schematic showing state transitions caused due to stochasticity. **B-D.** Stochastic simulation of model in Fig **3A** at 4 different volumes (0.01fL, 0.1fL, 1fL and 10fL from left to right) starting from the same initial concentrations and reaction rate parameters. Noticethe decrease in noise and reduction in the transition probability between fixed points. **E.** The stationary distribution of concentration-states for 100s simulations. Notethat in this model, the noise actually leads to an interesting effect where the time spent in the two stable-states changes asymmetrically with noise. **F.** Calculation of first passage time. The distribution P(X) of state vectors (X) was divided into 2 “stable-regions” by having distance cutoffs from deterministic stable fixed-points “0” and “1” (Methods). The transitions were calculated as jumps across these stable-regions. V(X) denotes the double-well potential shown in dotted lines, with 2 minima as the stable fixed points and T as the saddle point. **G.** A subset of topologies from which we could calculate residence time on 3 different volumes

Interestingly, we observed in a few cases that while the two peaks of the stochastic distributions were at the same position across volumes, the proportion in the first peak increased with volume (Figure 3E). In other words, the steady-state attractors of the bistable system were the same, but at small volumes, the system stayed longer near one fixed point, and at higher volumes, it shifted to the other fixed point. While we are not aware of previous reports of precisely this phenomenon, there are similar results for shifts of distribution using different kinds of noise on bistable dynamical systems (27). We calculated the transitions between the stable states, and hence the residence time distributions for all CRNs (Figure 3 F, G). As expected, the residence time and hence the noise robustness increases with volume, as the probability that the intrinsic noise causes state transitions reduces (28). The key finding here is that there is a wide range of noise robustness, even for networks measured at the same volume.

### Structurally robust systems contain many independently bistable subnetworks

We define Structural Robustness for a reaction system as the retention of the dynamical property of bistability with changes in the structure of the reaction system by removal of reactions. We found all network subsets of all 3561 bistable CRNs by removing one reaction at a time, as described in the Methods. We calculated the structural robustness as the fraction of subsets that were bistable. Our first observation was that structurally robust topologies were not preferentially enriched for any specific reaction type (Supp. Fig. S1).

(9) had shown that bistable dynamics run in CRN “families”, i.e., many bistable CRNs formed an interconnected set in a graph containing all configurations as nodes and addition (or removal) of reactions as edges. There were only 80/3561 (~2.2%) CRNs which could not be further broken down into smaller bistable CRNs. 56/80 of these minimal bistable CRNs were at the limit of the maximum network size we constructed, and hence we did not find larger CRNs that contained them. Consequently, the rest of the 3481 topologies originated from just 24 root motifs (Supp. Fig. S2). We called the set of topologies that were composed of a root CRN as the *root group* of the root CRN. We sorted and labeled these 24 root groups by the decreasing size of their family (Methods, Fig 4**B**, Table 1). For analysis of parametric robustness in the next section, we further filtered this group down to 8 root groups as over 95% (3350/3505) of the CRNs belonged to these 8 major root groups (Fig 4**B, C**). Group I, the most common root CRN is also the smallest bistable topology (Fig 1**C**, 4**B**) and was a subset of about 47% of bistable CRNs. Note that a single CRN can be a member of more than one root group. Similarly, many root CRNs could be constituents of a single, large CRN.

**Table 1.**
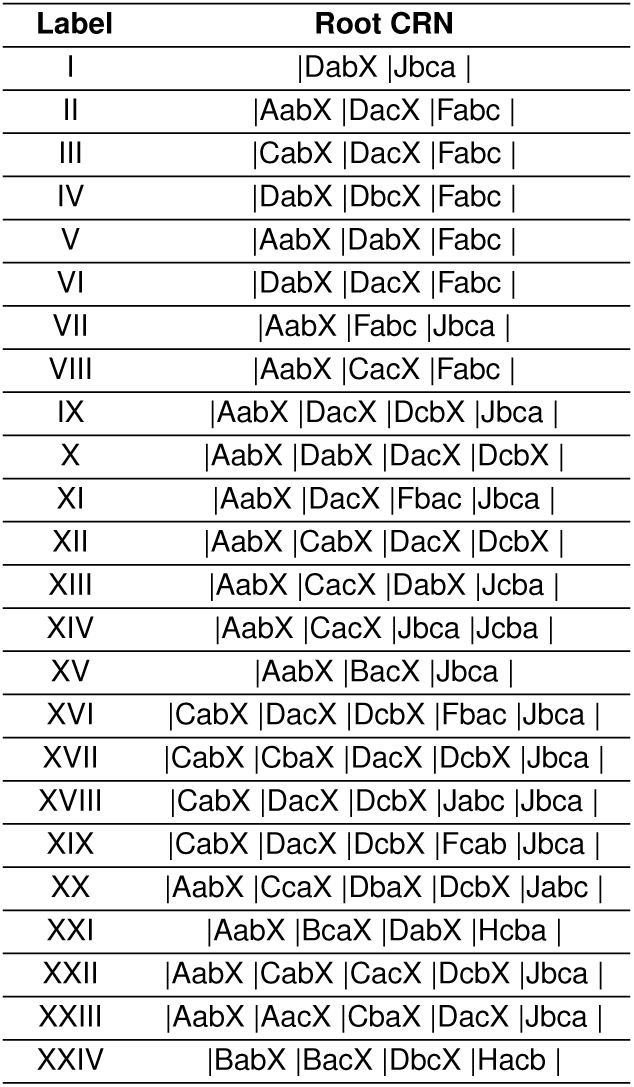
Root groups

**Fig 4.**
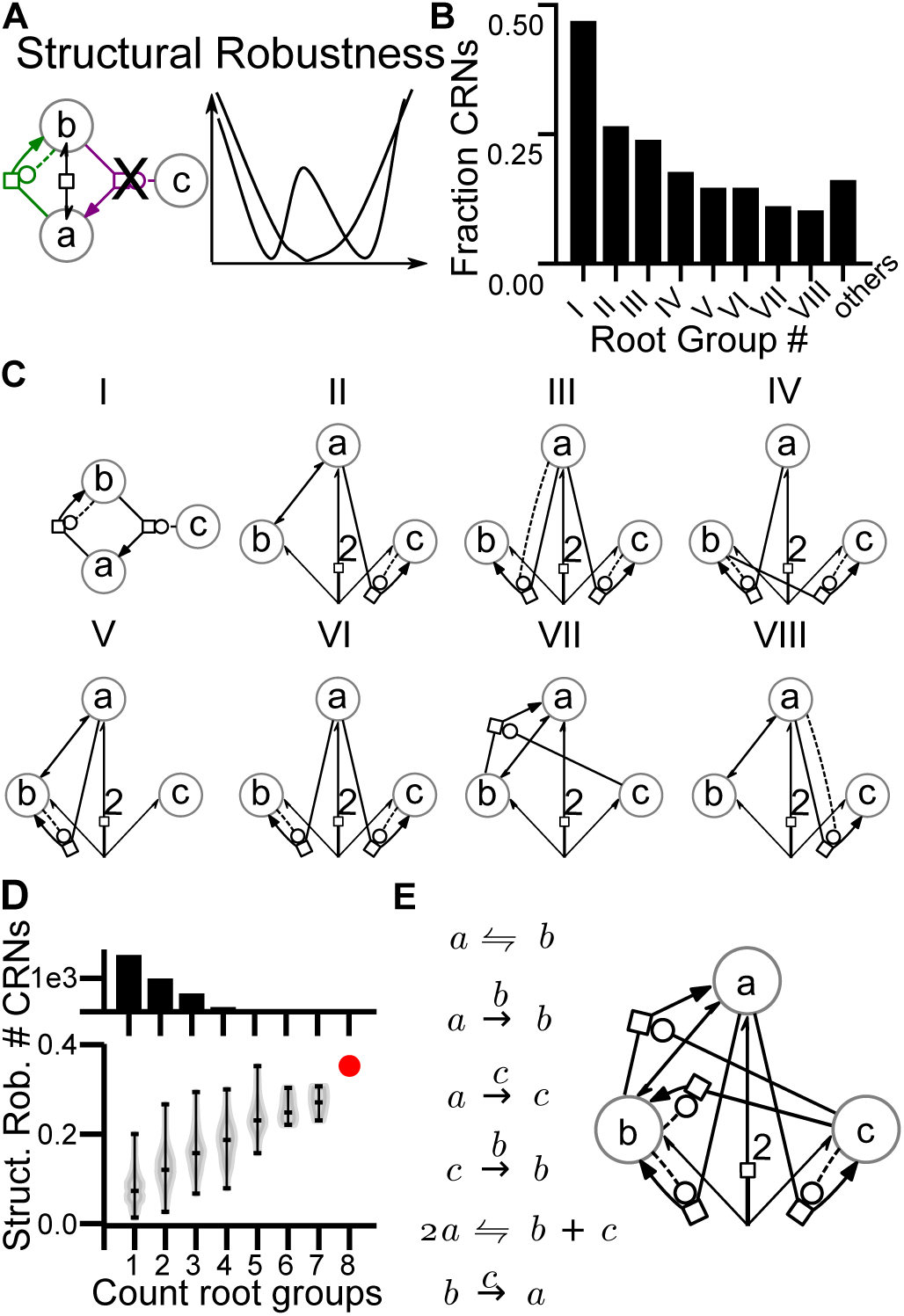
Structural robustness of bistable CRNs **A.** Schematic of structural robustness for example CRN showing loss of bistability due to removal of a chemical reaction from the CRN. **B.** A bar chart showing the fraction of total (3561) bistable CRNs that contain root CRNs as subsets. 8 out of 24 total root CRNs (as in **b.**) are shown here. The remaining 16 root CRNs were a subset of <5% of the total CRNs and have been shown grouped in "others”. **C.** Root CRNs as an SBGN graph for 8 out of 24 root groups, with group numbers on top in Roman numerals. **D.** Change in structural robustness of 3481 composite CRNs with an increase in the total number of root CRNs as subnetworks. Our highest structural robustness topology (marked in red) was also the only topology that was a member of 8 root groups. **E.** Most structurally robust topology in the database.

We found that members of root groups V and VI had the highest median structural robustness (Supp Fig S3). This implies that these root CRNs can grow via attachment of new reactions while retaining bistability. We then asked if structurally robust topologies contained multiple root bistables. We found that there was a strong correlation between the number of root bistables and structural robustness (Fig 4**D**). The structurally most robust CRN (Fig 4**D**) could be broken into 59 unique subset CRNs with 26 bistable and 8 root CRNs. In summary, composite bistable CRNs contained one or more of 24 root bistables, and the structural robustness of a CRN increased when it had more root groups.

### Parametric robustness was organized by root groups

We compared parametric robustness across the 8 largest root groups from Fig 4**B** and saw that root group VI had the highest median parametric robustness (0.77), followed by group III (0.74) and group IV (0.71) (Fig 5**A**). We found that out of the top 1% (35/3561) parametrically robust bistable topologies, 91.4% (32/35) had either the CRN in Fig 2**G** or Fig 5**E**. This explained the high symmetry observed in parametrically robust CRNs. These symmetric topologies were present in about 47% (167/356) of the CRNs in the top 10% (356/3561) of parametrically robust bistable CRNs. However, if we considered their frequency in all CRNs, they showed up only 16.3% (580/3561) of cases. Thus, having a pair of mutually-competitive autocatalytic feedback-loops yielded high robustness to parameter fluctuations.

**Fig 5.**
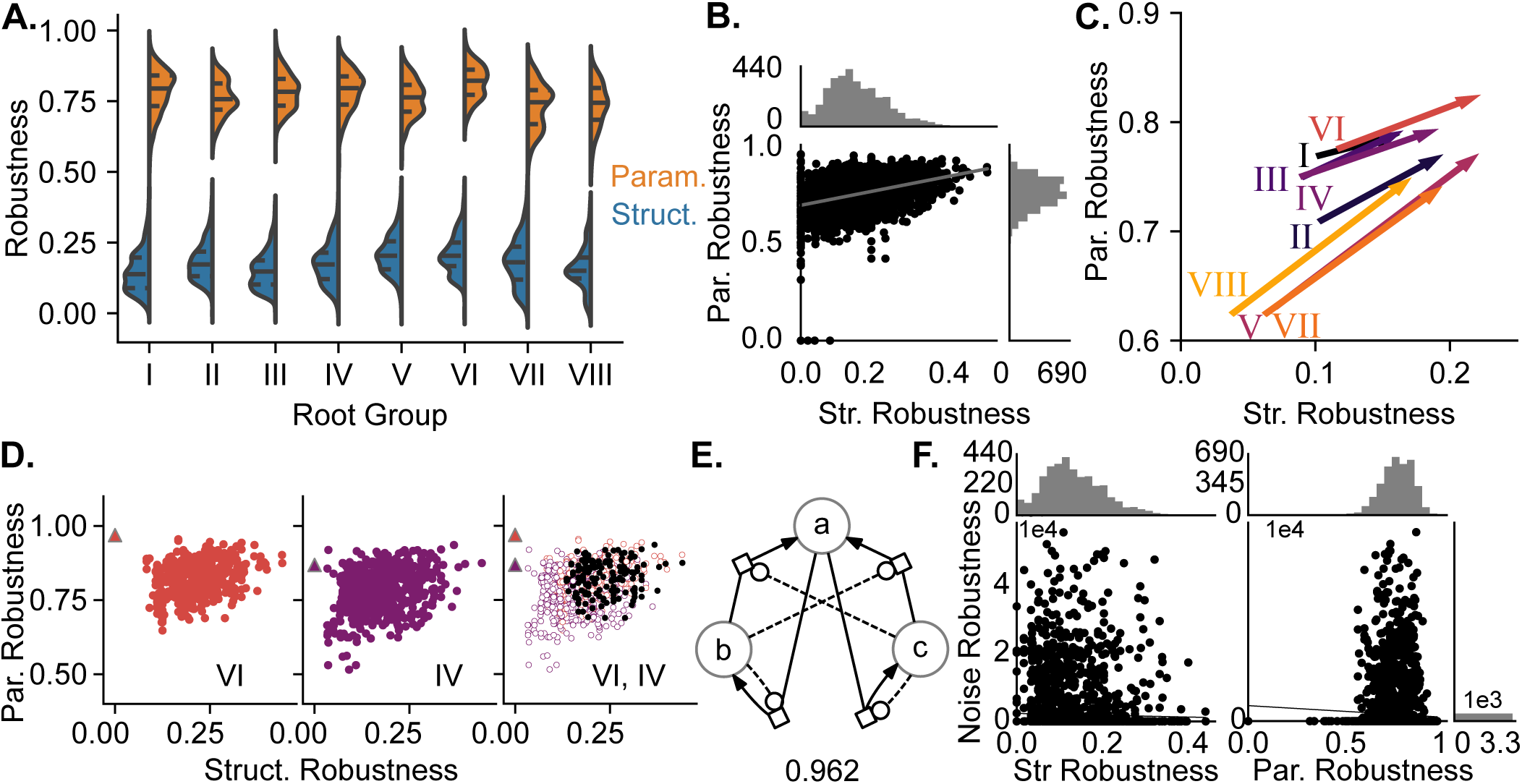
**A.** Distributions of structural (blue) and parametric robustness (orange) for CRNs grouped by their root topology from Fig 4**B**. Black lines inside the violin-plots mark the quartiles. **B.** Relationship between the structural robustness and parametric robustness for all 3561 CRNs with histograms showing their respective distributions. Slope of best fit = 0.43, R^2 = 0.12. All the root CRNs lay at 0 structural robustness. **C.** A comparison of the average structural and parametric robustness for CRNs belonging to each of the top 8 root groups. The start of the arrow represents the original robustness and the end o the arrow represents the robustness after the addition of other root groups. **D.** (Left) Relationship between structural and parametric robustness for CRNs belonging to root groups (Viand IV) in different colors. Colored triangles indicate the root CRN for the group. (Right) A pairwise overlay of the plots on the left, with CRNs that belonged to both groups colored black. **E.** Mirror-symmetric CRNs with coupled autocatalytic feedback loops showed high parametric robustness. **F.** Relationship of noise robustness with structural and parametric robustness.

A large fraction (44.5%) of members of root group VI had values of parametric robustness over 0.8.(Fig 5**A**). Remarkably, the root CRN of Group VI (Fig 2**G**) had a parametric robustness score of 0.97, that is, it was bistable for almost all 2000 randomly selected parameters. This CRN had a mirror-symmetric network topology. The root CRN of the third-highest parametric robustness (root group III) also shows a winner-take-all strategy (Fig 5**E**). In summary, membership of certain root groups, particularly group VI, conferred high parametric robustness on CRNs.

### Correlation between structural, parametric, and noise robustness

Next, we asked if topologies that are robust to structural perturbations also show robustness to parametric perturbations (Fig 5**B**). We found that the values for these forms of robustness were indeed correlated (Pearson’s correlation coefficient = 0.35). This has the surprising implication that interlinking multiple independently bistable root groups added to the range of parameter space in which these CRNs retained bistability, instead of interfering with each other’s parameter space to destroy bistable behavior.

Surprisingly, almost all of the very high structural and high parametric robustness combination CRNs were members of root group VI (Fig 5**B-D**). When we checked for CRNs which had only root group VI, we found that they still had high structural and parametric robustness, which shows that this group is generally robust to the addition of reactions, has flexibility in its parameter space and gregariously combines with other root CRNs to create composite robust bistable CRNs Fig 5**C**.

We also compared the parametric and noise robustness for all CRNs where we had all three measurements (Fig 5**F**). We found that structural and parametric robustness had no average correlation with noise robustness.

### Related CRNs have similar structural and parametric robustness

We then looked at the relationship between structural and parametric robustness by grouping the data with respect to the two of the most parametrically robust groups: IV and VI (Fig 4**C**, 5**D**). In Fig 5**D**, where we show topologies containing either group VI (left, orange) or group IV (middle, purple) or a combination of both (right, black). It can be seen that the combination (black) has high parametric robustness, closer in value to group VI than to group IV. In addition, by virtue of membership of two (or more) root groups the structural robustness of the CRN also increased leading to a topright placement of composite CRNs (Fig 5**C,D**). Thus, CRNs which were members of group VI root, in addition to other root groups were more likely to be both structurally and parametrically robust CRNs.

## Discussion

We have compared different robustness properties of 3561 bistable CRNs in the SWITCHES database with ≤ 6 reactions between ≤ 3 molecules and 3 reactions between 4 molecules. We find that robustness to noise does not correlate with either structural or parametric robustness, but there is a small positive correlation between the latter two. This arises substantially because of a ‘family dependence’, as follows: 3505/3561 (98.4%) of bistable CRNs were derived from one of 24 root bistable CRNs, and descendants of some of these root CRNs had consistently higher structural and parameter robustness.

### Generalizing robustness

Bistable systems offer a clean way to test for the robustness of function, through dynamical systems analysis yielding a yes/no result for the presence of the function (29), and large sample size (9). Thus we have been able to explore three distinct and well-defined dimensions of robustness in chemical bistables. Do these results carry over to other systems?

A simple approach to generalizing is to define a functional range for a systems property, such as amplification. The system is functional only if its gain is within this range. With this binary classification, we can apply similar measures for robustness as in the current paper, for the dimensions of parameters and topology. Such estimates of robustness have frequently been made in the literature. Parameter robustness is commonly estimated through parameter sensitivity analyses in signaling (17, 30). Topology robustness has been extensively analyzed in terms of reaction motifs and their role in systems function (17). In parallel with our observation that certain ‘families’ of bistable topologies may be particularly parameter-robust, there are several suggestions that certain core reaction motifs may also confer parameter robustness (17, 31). For example, inhibitory feedback tends to linearize system output and make it less dependent on rate terms (32). Thus we suggest that non-bistable signaling functions may also exhibit weak correlations between parameter and topology robustness, arising from the presence of core reaction motifs.

While it is more speculative to extrapolate outside the chemical domain, we do note that many natural and engineered systems fall in dynamical families similar to chemical networks. For example, the Lotka-Volterra population model is formally similar to a chemical oscillatory model (33). Further, there is a formal similarity between flux chemical networks, and nutrient flow through food webs (34). A key limitation in such analogies is that we have sampled randomly, and exhaustively, whereas nature does not. For example, the ‘nodes’ in a food web are species with distinct evolutionary histories that include special survival mechanisms and responses to extreme conditions. Our conclusions about robustness do not factor in such selection bias.

### Selection for motifs determined by both function and robustness

Network motifs, or subnetworks that occur more frequently than expected by chance, are seen as an important structural organizing feature of real biological networks (35). Since motifs can perform fundamental nonlinear computations such as bistability, gain control, and oscillations, they were also suggested as the reusable functional units of complex networks (36, 37). The idea of evolutionary selection based on motif function has been criticized due to the non-unique mapping from network structure to function (38). Around the same time, it was shown that the abundance of motifs is directly correlated with the robustness of motifs to perturbations (7). We find that reaction systems with the same function (bistability) have different levels of robustness to different perturbations. Thus, even if robustness is a basis Str Robustness Par. Robustness for motif selection, two functionally equivalent motifs may fare very differently under different conditions. One potentially simplifying factor is our observation of a weak correlation between two forms of robustness for parameter change and network topology change (mutations). Thus, even though there is a degeneracy in network structure, robustness to perturbations may end up selecting networks with a similar structure. There is some degree of support for this idea. Specifically, we find that the highly robust root CRN group VI, is mirror-symmetric. Here, a molecule can catalyze its formation, as well as inhibit its inhibitor to lead to double-positive feedback. Certain mirror-symmetric structures have been reported earlier in different systems and have been termed *reciprocal regulation* or *cross-inhibition*, e.g., the CDK1-Cdc25-Wee1 system for mitotic cell-cycle transitions (39), cellular motility by chemoattractant sensing (40), and the GATA-1 - PU1 for cell-fate determination in multipotent progenitor cells (41).

#### Robustness and the evolution of enantiomeric asymmetry in biomolecules

Robust bistability is also relevant for enantioselective synthesis in biological systems. Bistable systems have long been proposed as a chemical means of abiogenic selection of L-type or D-type enantiomers observed in biology (42). Parametrically robust enantioselective mechanisms can sustain and allow for the creation of a variety of different biomolecules by being resilient to substrate and enzyme alterations without losing bistability. Curiously, the group VI root CRN (Fig 2**G**) is similar to the chemistry proposed by (13) for the synthesis of pure enantiomeric (only L or only D-forms) from an achiral substrate and later experimentally validated by the Soai reaction (43).

This mechanism can be understood by considering *a* as the achiral precursor of the pure enantiomeric forms (*b* and *c*) of the product, which catalyze their own formation (Fig. 2**G**). In this CRN, *b* and *c* react together to give back 2 molecules of *a*. Note that the reaction network suggested for the generation of homochirality shows up as the top parametrically robust reaction system (group VI), and this CRN is also robust to the addition of new reactions. Thus, once this reaction topology has evolved for the asymmetric synthesis of enantiomers, it is conducive to the addition of further reactions to increase efficiency, without losing its bistable dynamics. (14) proposed an alternative model with cross-catalysis to generate back *a*, which surprisingly, shows up as our second highest parametrically robust topology shown in Fig. 5**E**. It is worth noting that this CRN is a mirror-duplication of our smallest possible bistable CRN across the axis through *a* (Fig 1**D**)

In summary, our exploration of robustness using bistable CRNs suggests that for key categories of dynamical systems, the robustness of different types is mostly uncorrelated. The presence of ‘family advantages’ for certain network topologies may steer evolutionary trajectories in cases where robustness provides a selective advantage.

## Methods

### Bistable CRN selection

Bistable CRNs were extracted from Searchable Web Interface for Topological CHEmical Switches (SWITCHES, http://switches.ncbs.res.in), an online resource curated using the results of (9). SWITCHES contains ~7e6 chemical kinetic models of 2-5 reactants with 35967 bistable models in cspace and SBML formats along with their steady states. **3561** bistable CRNs were extracted from these bistable models. This is an exhaustive list of all bistable CRNs detected in the original survey, with up to 3 reactants and 6 reactions or 4 reactants and 3 reactions.

### Graph Generation

SBGN graphs (44) were generated for all CRNs using elementary chemical reactions depicted in Figure 1**a**. We used Python with the free graph visualization library Graphviz (45) and implemented a Cspace CRN graphing library to generate SBGN graphs.

### Simulations

All simulations were performed in Python 2.7 the Multiscale Object Oriented Simulation Environment (MOOSE) (46). The deterministic steady-state finding routine uses deterministic simulations of the coupled chemical reaction differential equations based on root finder using GSL (47) and extraction of conservation laws from the stoichiometry matrix (48). The stochastic simulation was also performed in MOOSE, which uses the Gillespie algorithm (49). Steady State Solving and Stochastic simulations were done on 1000 cores in an array job on a cluster computing framework using Sun Grid Engine (SGE), bash and Python.

#### Generating random models

Bistable CRNs selected from the SWITCHES database were populated with rates logarithmically sampled over 6 orders of magnitude. Since we typically had to sample over a large number of parameters (4 to 12), an efficient sampling scheme was utilized to sample the rates efficiently. We implemented an efficient sampling method called Latin Hypercube Sampling with multi-dimensional uniformity (23, 24). In this method, the parameter space is divided into strata by oversampling (5 times) using a uniform sampling method. Then these grids are sorted, and the pairwise distances are calculated between all pairs of strata. This is followed by calculating the average distance of each stratum with its neighbouring stratum. Finally, the stratum with the minimum average distance from other strata is eliminated. This process is repeated until the required number of samples is reached. Real numbers between consecutive strata are then sampled uniformly using conventional Monte Carlo methods. This helps in maintaining reasonable uniformity of sampling in high dimensional space that is not achievable using simple Monte Carlo methods. This implementation is publicly available at https://github.com/sahilm89/lhsmdu. More details about the algorithm can be found in (23). In this way, we sampled 2000 sets of random rates, with two rates for each reaction in the CRN. For reversible reactions, this was forward (*k*_1_) and backward (*k*_−1_) rates. For enzymatic reactions, although we sampled *k_m_* and *k_cat_* from the Michaelis Menten formalism, they were internally converted in MOOSE into mass-action formalism, with 4*k*_−1_ = *k*_2_ = 4*k_cat_* and 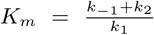.

#### Finding steady states

We used the steady-state solver in MOOSE to find steady-states for these models, which uses time-course analysis in conjunction with linearization around the fixed point to find and assess the dynamical behavior of steady-states. We systematically swept across the initial condition space by random, stoichiometrically-constrained jumps and found steady states. Then the algorithm analyzed the stability of the fixed points found based on the signs of the eigenvalues of the Jacobian matrix. While this process does not guarantee that all possible fixed points will be found, it is faster than other algorithms, e.g., homotopy continuation (9). We started from 1000 different initial conditions to get a list of steady states for each model. This process was repeated for all 2000 random rates for the topology.

#### Classifying bistables

We clustered the steady states obtained from above into one or points to classify a model as bistable if there were exactly two clusters found. We used to the following elementwise distance measure for this classification:

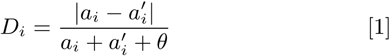

*a_i_* is the concentration of the entity at the *i^th^* index in the steady-state vector being tested for similarity against 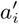, the corresponding concentration in the running-average solution vector(s), *θ* is the offset = 1e-3, and where *D_i_* is the element-wise distance of the *i^th^* entity. The offset *θ* avoids divisions by zero. If *all* the elements of the solution vector fall under 5% tolerance, the vectors are considered identical. If this process yields two sets of solutions at the end, the model is called bistable.

### Structural Robustness

We quantified Structural Robustness (*S*) as the number of subnetworks of the CRN that were bistable normalized to the total number of subnetworks.

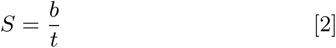

where b is the total number of subnetworks that are bistable, and *t* is the total number of subnetworks in the topology. We utilized the string-based string-based (Cspace) format of the CRN for counting subnetworks, as explained below.

### Subnetwork counting

To count the subnetworks, we first took the reaction signatures for the CRN and created all possible combinations of the reactions, keeping the relative connections between reactants constant. For example, the CRN from Fig 4**b**, |*DabX*|*DacX*|*Fabc*| would have the following combinatorial subnetworks: {|*DabX*|, |*DacX*|, |*Fabc*|, |*DabX*|*DacX*|, |*DabX*|*Fabc*|, |*DacX*|*Fabc*|}. Also, for some reversible reactions, certain node labels can be permuted without loss of relative relations between reactants. For example, reactants *a* and *b* can be permuted in signature |*AabX*| to give|*AbaX*| without changing the meaning of the reaction. This is also true for permutations of molecules *b* and *c* in reactions |*Eabc*|, |*Fabc*| and |*Kabc*|. We treated these permutations equivalently at the second step to generate the set of all subnetworks. Lastly, the node labels were globally permuted, i.e., all possible permutations of the reactant label set {*a, b, c, d*} were done to generate the final subnetwork set for each CRN. Then, for each topology, we compared these subnetworks against bistable CRNs and counted the exact matches.

### Symmetry analysis

The node label permutations mentioned above were also used for symmetry analysis. We looked for independent subnetworks for a given CRN that were identical to each other. If these subnetworks were sufficient to construct the entire CRN, the CRN was called globally symmetric. 36 such CRNs were found. If other reactions besides the identical independent subnetworks were required to constitute the CRN, the network was called locally symmetric, and the number of reactions that needed to be added was called the order of the CRN. Yet other CRNs had no symmetry at all.

### Parametric Robustness

Parametric Robustness, *P* was quantified as the fraction of the n-dimensional volume of the bistable parameter space. It was calculated as the fraction of the total number of random parameter models for a given topology that were bistable normalized by raising to the power of 1/number of parameters.

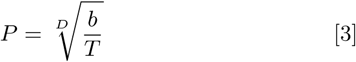

where *b* = Number of bistable parameter sets, *T* = Total number parameter sets, and *D* = Number of parameters in CRN.

### Noise Robustness

Noise robustness for a bistable model at a given volume was quantified as the geometric mean of the average time spent in the basins of the two stable fixed points before the system switches to the other fixed point due to statistical fluctuations. The time spent before the transition in each state is called the Residence Time (RT) (50), or Kramers’ time (51). For a CRN, the Noise robustness was averaged over all parametrized models.

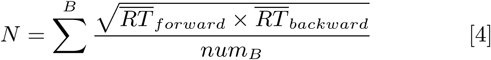

where, *B* = Across all bistable models

*num_B_* = Number of bistable models

*RT* = Residence Time

Forward and Backward Kramers’ times 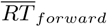 and 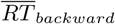 for a given volume were calculated by starting off a stochastic simulation using Gillespie’s exact stochastic simulation algorithm (49) and measuring time before a state transition. We measured transitions using a cutoff on the continuously calculated element-wise distance between the measured stochastic state vector, and two deterministic fixed points using Equation 1 with *θ* = 1e-1. This was then repeated for various volumes from 1e-21 *m*^3^ to 1.024 e-18 *m*^3^ in 10 logarithmic steps of base 2, with a simulation time of 1 day (86400 seconds), or until a spontaneous transition happened. These volumes were chosen as they are in the range of typical small compartments such as dendritic spines, but interestingly, it also happens to be the parameter space where stochastic effects are relevant in the chemical reaction networks that have realistic reaction-rate parameters. The whole run was repeated 50 times.

### Removing stuck states

In some CRNs with irreversible reactions, all gradients and all but one concentration can become zero, called a “stuck” state. Once the system achieves this state, it is not possible to escape from it with intrinsic fluctuations, because all transition probabilities are zero which leads to a no-noise state. Thus, before running stochastic simulations, the continuous fixed point vectors (in concentrations units) were converted into discrete molecule counts at 100 *fL*. If there were models whose steady states had less than 2 non-zero elements in their state vector, they were labelled “stuck” and were omitted from stochastic simulation and further analysis.

## Supporting information

Suppmentary Figures

## Acknowledgments

We would like to thank Sandeep Krishna for numerous suggestions over the development of this project, and Avrama Blackwell for comments on a draft version. We thank CSIR, NCBS-TIFR, DAE, and DBT (BT/PR12255/MED/122/8/2016) for funding. Large calculations were performed on the campus Supercomputing Facility supported by NCBS-TIFR.

