## Supplementary material for "Different dimensions of robustness - noise, topology and rates - are nearly independent in chemical switches": Suppmentary Figures

### Supplementary Figures

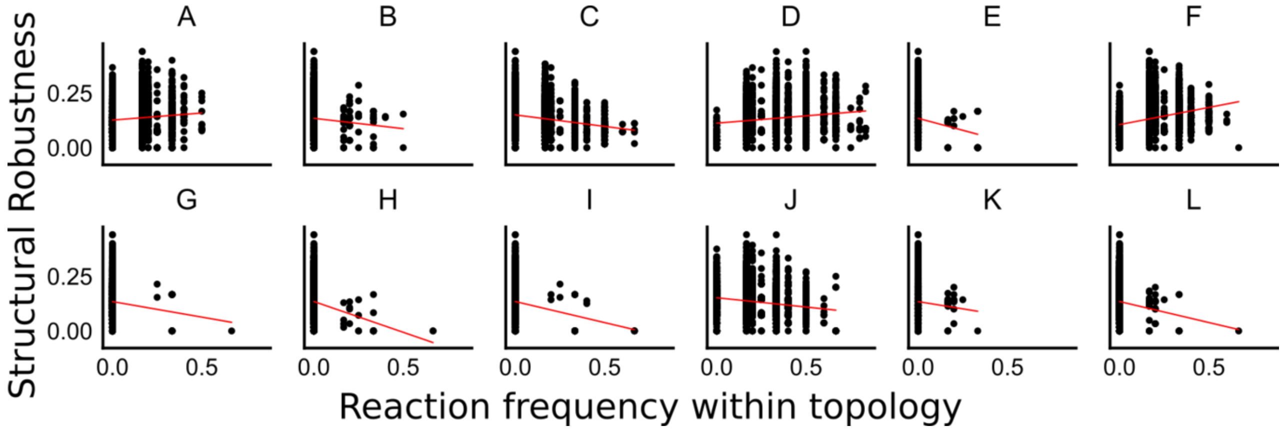


***S1.*** *The relationship between the Structural Robustness for a topology, with the fraction of occurrence of a given reaction type within the topology. None of the reactions shows a strong correlation with structural robustness, suggesting that different structures may be required for different topologies to increase Structural Robustness.*

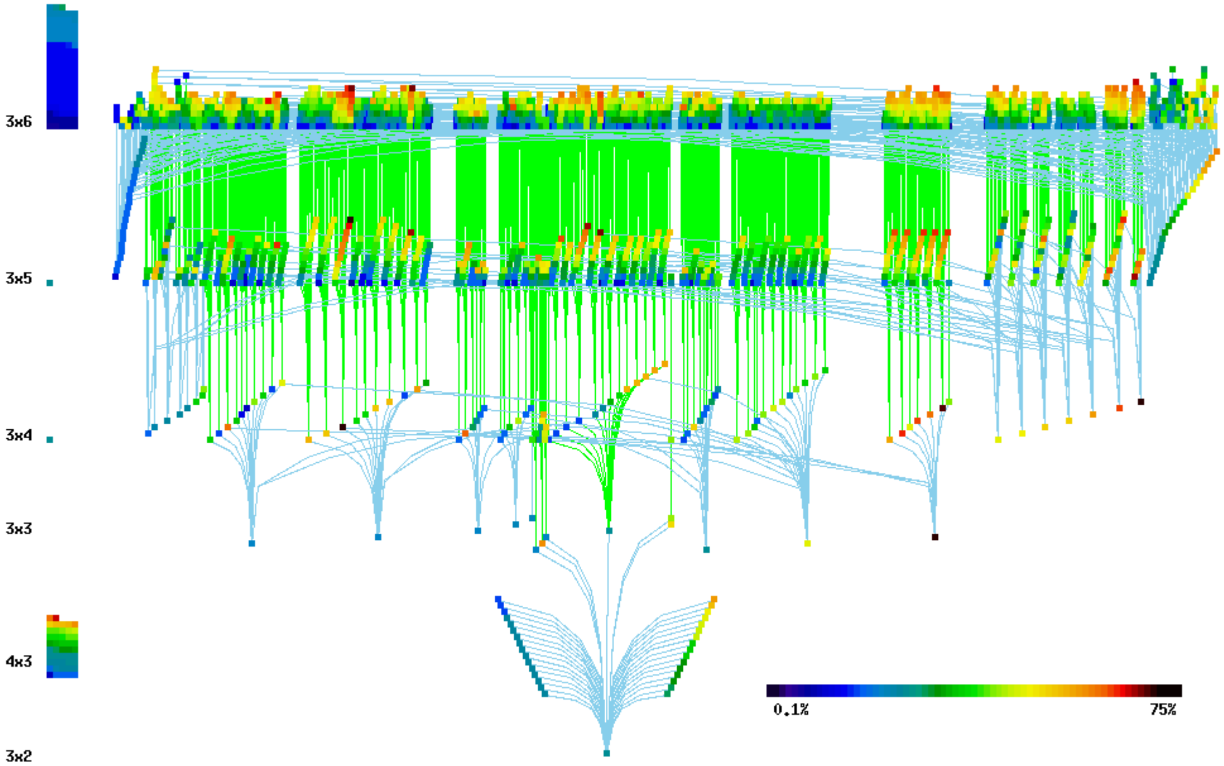


***S2**. Taken from Ramakrishnan and Bhalla (2008, doi: https://doi.org/10.1371/journal.pcbi.1000122). This figure shows the relatedness (edges) between the different CRNs (nodes) and their propensity to be bistable (node colors). Different layers denote different dimensionalities (labelled on the left) of the CRNs. Nodes without any edges coming from bottom (smaller dimensionality) are the 80 root CRNs. This includes both 56 orphan nodes (stacked on the left), and the 24 root group CRNs.*

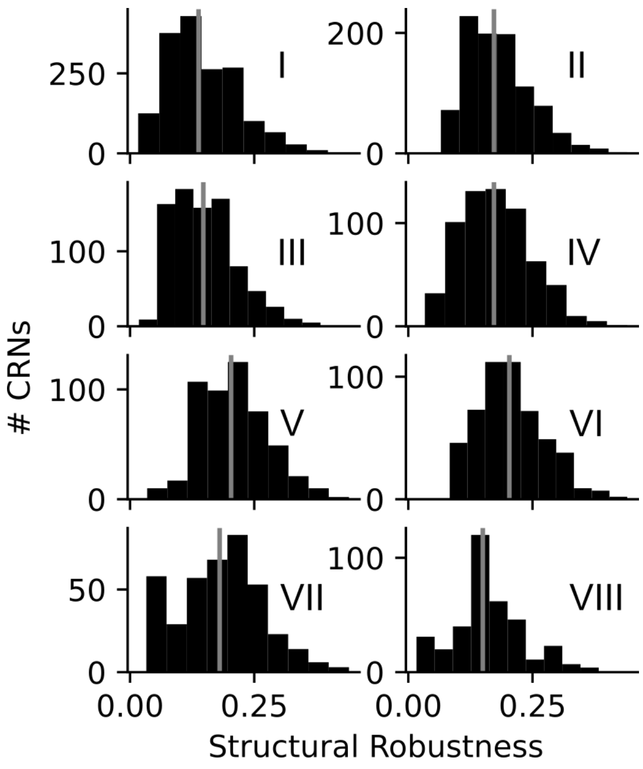


***S3**. Distribution of structural robustness of CRNs organized by root groups. Group V and VI have the highest median structural robustness followed by Group VII. Group V and VI had about 30% members common to both the root groups.*
